# Mutation hot spots for clinical pathogenicity across the SLC6 transporter family

**DOI:** 10.1101/2025.01.24.634481

**Authors:** Jiahui Huang, Daniela Digles, Gerhard F. Ecker

**Affiliations:** University of Vienna, Department of Pharmaceutical Sciences, Vienna, Austria

## Abstract

Genetic mutations of the Solute Carrier 6 (SLC6) family can lead to a diversity of clinal syndromes, such as creatine deficiency. Studying the impact of genetic mutations at the SLC6 family level is valuable not only for their medical significance but also for their conserved sequence and structural features. Within this work, we aim to link the disease-related mutations to their clinical significance from protein-protein interactions (PPIs) perspective, addressing how particular mutations may affect these critical interaction hotspots.

In this study, we integrated both curated mutation data from previous work and predictive output from AlphaMissense to examine the entire SLC6 family. The mutations were mapped both onto the sequences and structures of SLC6 transporters. Thereby, a clustering of pathogenic mutations appeared on the surface regions that are likely involved in PPIs. After modeling complexes of SLC6s with potential shared interactors, we assessed these models overall and interface quality. By analyzing the complex interfaces together with the pathogenic mutations, we identified specific hotspots in the interfaces enriched with pathogenic mutations. In-depth examinations of selected PPIs offered insights into how particular mutations may affect these critical interaction hotspots.

The hotspots were identified on the ECL3 and ECL4. For instance, Thr394Lys in SLC6A8 was found in the model interfaces, supported by experimental data showing the significant enrichment of the mutated proteins in the ER. Understanding these mutation hotspots can shed light on the broader structure-function relationships of SLC6 transporters and encourage therapeutic interventions targeting protein-protein interactions affected by pathogenic mutations.

## Introduction

The Solute carrier (SLC) 6 transporter family, which includes 19 genes (SLC6A1-SLC6A20) and one pseudogene (SLC6A10), is a critical component in cellular transport and regulation. These transporters can be categorized into four subgroups based on sequence similarity and substrate specificity: monoamine neurotransmitter transporters (SLC6A2, SLC6A3, SLC6A4), amino acid neurotransmitter transporters (SLC6A5, SLC6AA7, SLC6A9, SLC6A14), nutrient amino acid transporters (SLC6A15-SLC6A20), and the other amino acids transporters (SLC6A1, SLC6A6, SLC6A8, SLC10-SLC6A13) ^1, 2^. The first two subgroups, known as the neurotransmitter transporters (NTTs), include nine transporters involved in neurotransmitter uptake and regulation, which are essential for central nervous system (CNS) ^3^ signaling.

Mutations in SLC6 transporters can affect their function by either directly disrupting their transport activity or indirectly affecting the protein-protein interactions (PPIs). Disruption of these interaction sites can compromise the transporter’s integration into essential cellular processes, contributing to disease development. These mutations have been linked to a variety of Mendelian disorders ^4–6^ and complex multifactorial diseases ^7, 8^, highlighting the importance of understanding complex indirect pathogenic mechanisms of SLC6 mutations.

An estimated number of PPIs in human cells – termed as interactome - was provided by Lu et al., which ranges from 130,000 to 650,000 ^9, 10^. Yet, the number of available protein complex structures is far less. To bridge this gap, computational approaches like protein-protein docking and structural modeling are often used. Protein-protein docking, adapted from ligand docking, assumes the relative rigidity of individual protein structures, predicting interactions based on surface adjustments^11^. In contrast, structural modeling approaches treats PPIs as dynamic, flexible complexes, offering a more comprehensive view by integrating the entire complex rather than focusing solely on the interface. Another difference between protein-protein docking and structural modeling approaches is that pre-solved structures of individual proteins are prerequisites for the former but not the latter.

Recent advancements, particularly in deep learning models like AlphaFold ^12^ and AlphaFold Multimer (AF-M) ^13^, have leveraged our ability in complex protein interaction predictions. AlphaFold Multimer, optimized for modeling protein-protein interactions ^14, 15^, is well-suited for the investigation of SLC6 transporter interactions. Using data from sources like the RESOLUTE database^16^, which provides specific PPI information for SLC6 family members, we can identify shared interaction partners and gain insights into the regulatory networks these transporters participate in.

In our previous work ^17^, we collected a set of mutations with curated clinical significance annotated by binary labels (pathogenic or benign). A small fraction of this dataset was tested for their expression, location, and function. Building on this, our current study examines these mutations from a structural perspective, seeking common patterns among pathogenic mutations. Our findings suggest that cluster of pathogenic mutations on the shared surface region of SLC6s can play critical roles in PPIs, providing a potential explanation for their effects across the SLC6 family. Additionally, this work provides the research community with the following tools and resources: First, a KNIME tool which allows to visualize the location of their mutations of interest in a 3D structure. Second, multiple scripts fine-tuned for SLC6s PPI network analysis and AF-M structure analysis. Third, a rationalization of the potential impact of specific pathogenic mutations on the SLC6 PPI interfaces.

## Results

### Sequence alignment and structure superposition

19 complete canonical sequences were retrieved from UniProt for the whole human SLC6s, excluding the pseudogene SLC6A10. An alignment file was generated utilizing the Clustal Omega tool integrated in the EMBL-EBI service. With the help of MOE, the alignment file was loaded and interpreted with a pairwise similarity matrix. Viewing from the generated matrix (Fig. 1), this alignment revealed a conserved pattern consistent with the current knowledge about the SLC6 family sequence ^18^. The resulting full sequence pairwise similarity spanned between 36% to 81%.

**Fig. 1.**
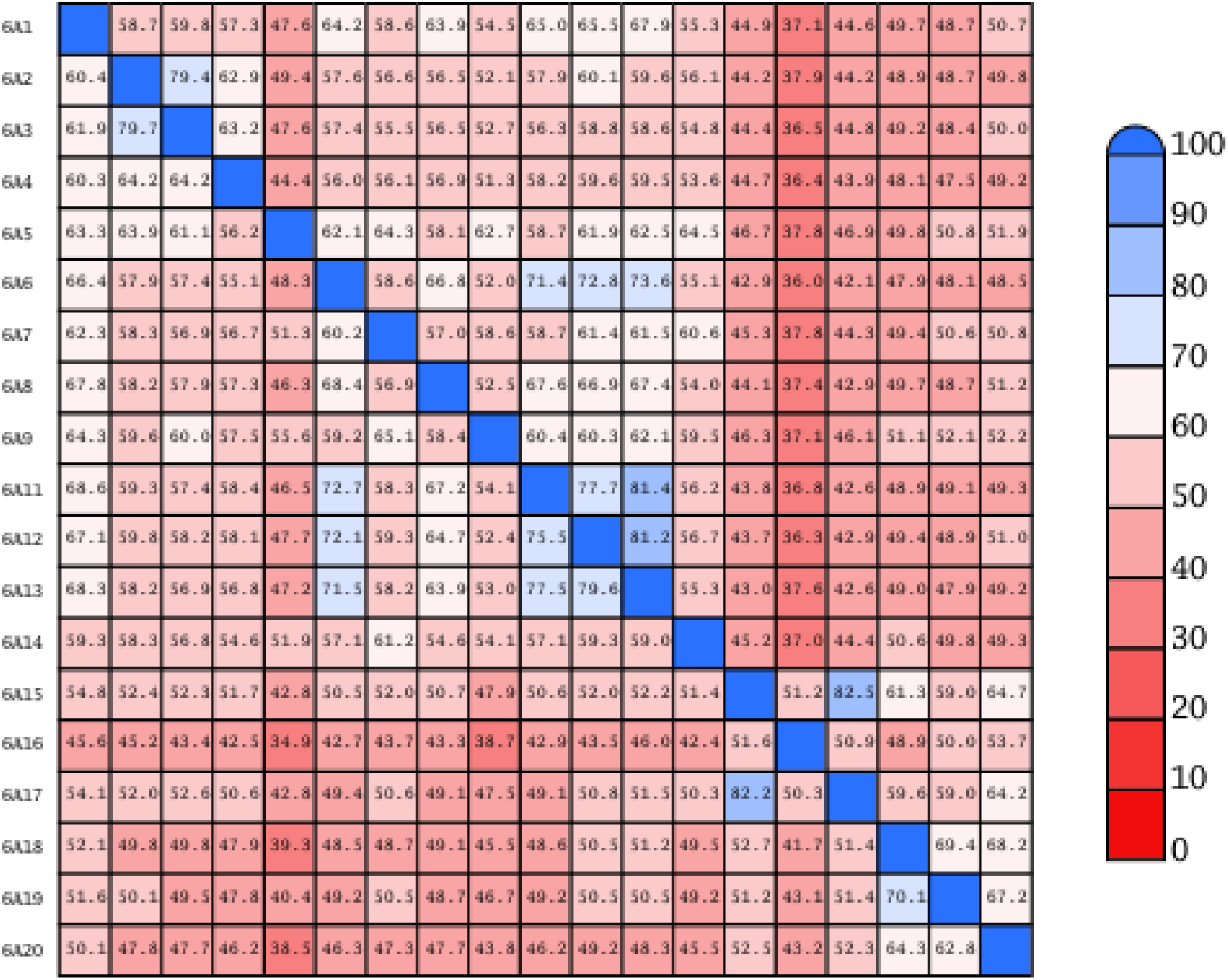
Heatmap of the pairwise similarity matrix of SLC6 canonical sequences. The sequence is ordered numerically based on the gene name.

Besides the adequate sequence similarity, the SLC6 family is defined with a conserved structural pattern. This pattern is known as the LeuT fold, characterized by 12 transmembrane α-helical domains (TMs 1-12) linked by intracellular loops (ICLs) and extracellular loops (ECLs). As not all the structure compositions were solved with the limited available structures published in the Protein Data Bank archive (PDB), full sequence structure predictions for all 19 SLC6s were retrieved from the AlphaFold Protein Structure Database (AlphaFold DB version 4) ^19^. Based on the imported sequence alignment, 19 predicted structures were superposed (Fig. 2). In addition to the default superposition setting using the current alignment, two other weighting options are selected to achieve a better view of the conserved structural pattern of this family. The first was to enable “Accent Secondary Structure Matches” to leverage the weight of matching defined secondary structure. The second was to turn on the “Refine with Gaussian Distance Weights” function, which can improve the superposition with an iterative algorithm with dynamic residue weights ^20^.

**Fig. 2.**
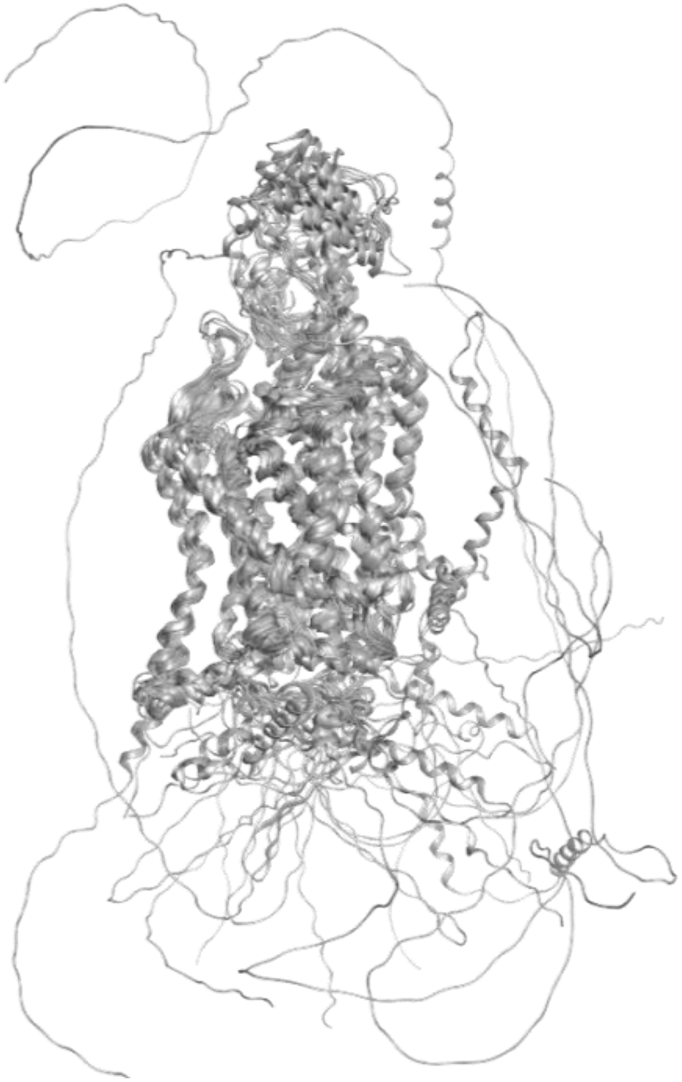
Superposition of SLC6s structural models. The models are extracted from the AlphaFold Protein Structure Database (V4). The superposition was performed by prioritizing the defined secondary structures that correspond to the TM regions.

Note that the AlphaFold predictions are presented as full-sequence structural solutions. They include flexible loops and certain unfolded regions, which make the superposition less tidy. This can hardly be avoided without sacrificing the completeness of the structure. Nonetheless, due to a higher weight of the defined secondary structure and the increased dynamic iteration of the superposition process, the transmembrane region of the structures was presented in a well-overlaid mode with an overall CA-weighted RMSD (wRMSD) value under 2 Angstrom (Fig. 3).

**Fig. 3.**
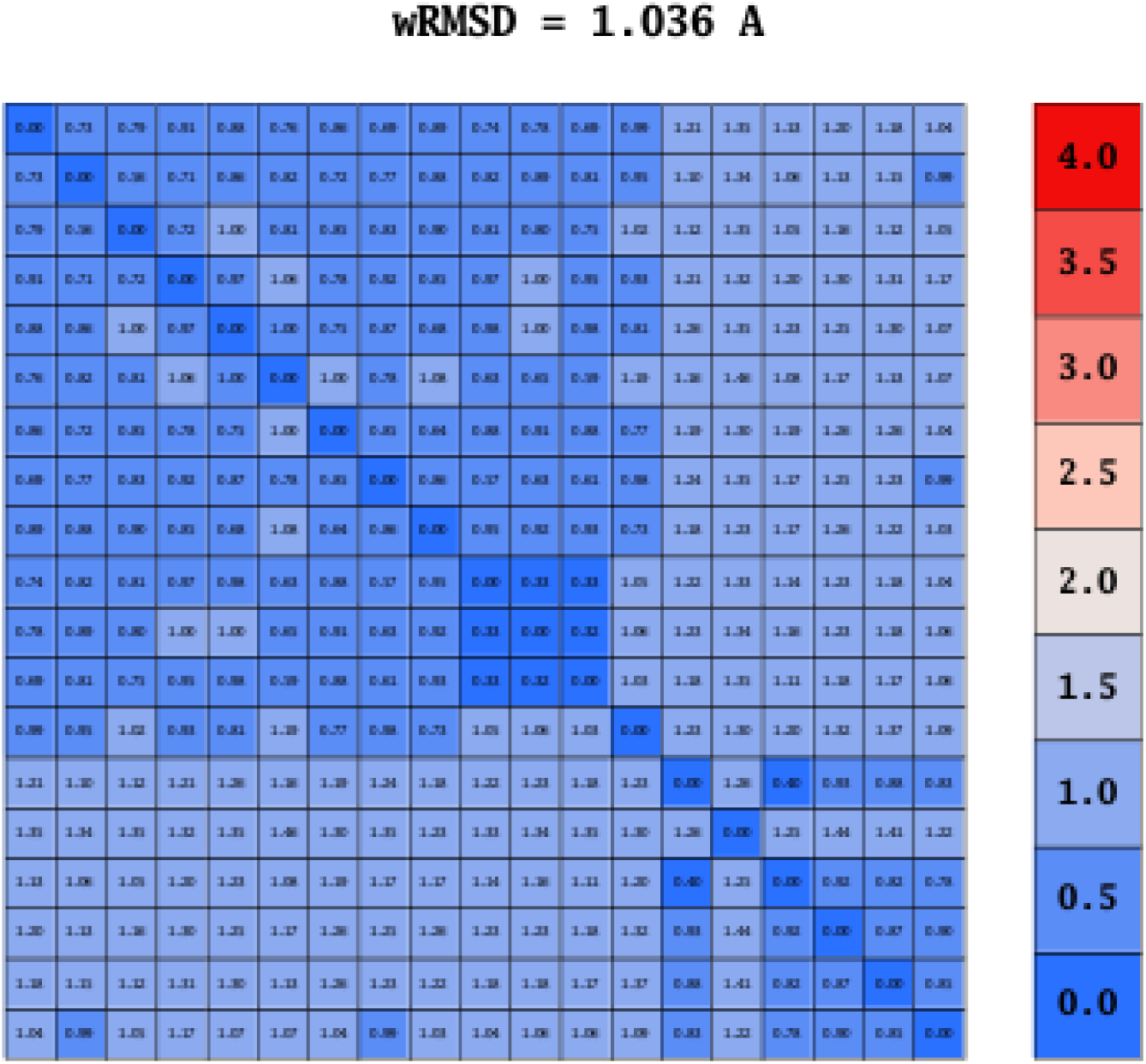
Heatmap of pairwise weighted RMSD (wRMSD) of the transmembrane region CA atoms of SLC6 structural models extracted from the AlphaFold Protein Structure Database (V4). A higher weight was signed to the defined secondary structure in an iterative structure superposition process.

### Sequence logos

A sequence logo was generated from the aforementioned alignment file with the help of the Seq2logo web server. This presentation of the alignment can provide not only information about the stack height for residue probability but also the stack width for the occupancy of the aligned position. An example of the TM6 is shown in Fig. 4A. The highly conserved Gln and Gly (position on the alignment: 568 and 578) are reported to be crucial for the function of multiple SLC6s. Gln’s role in the transporter function can be chased back to its location in the chloride ion-binding site ^21^. Gly was reported to be an essential determinant of protein dimerization and cell surface expression ^22^.

**Fig. 4.**
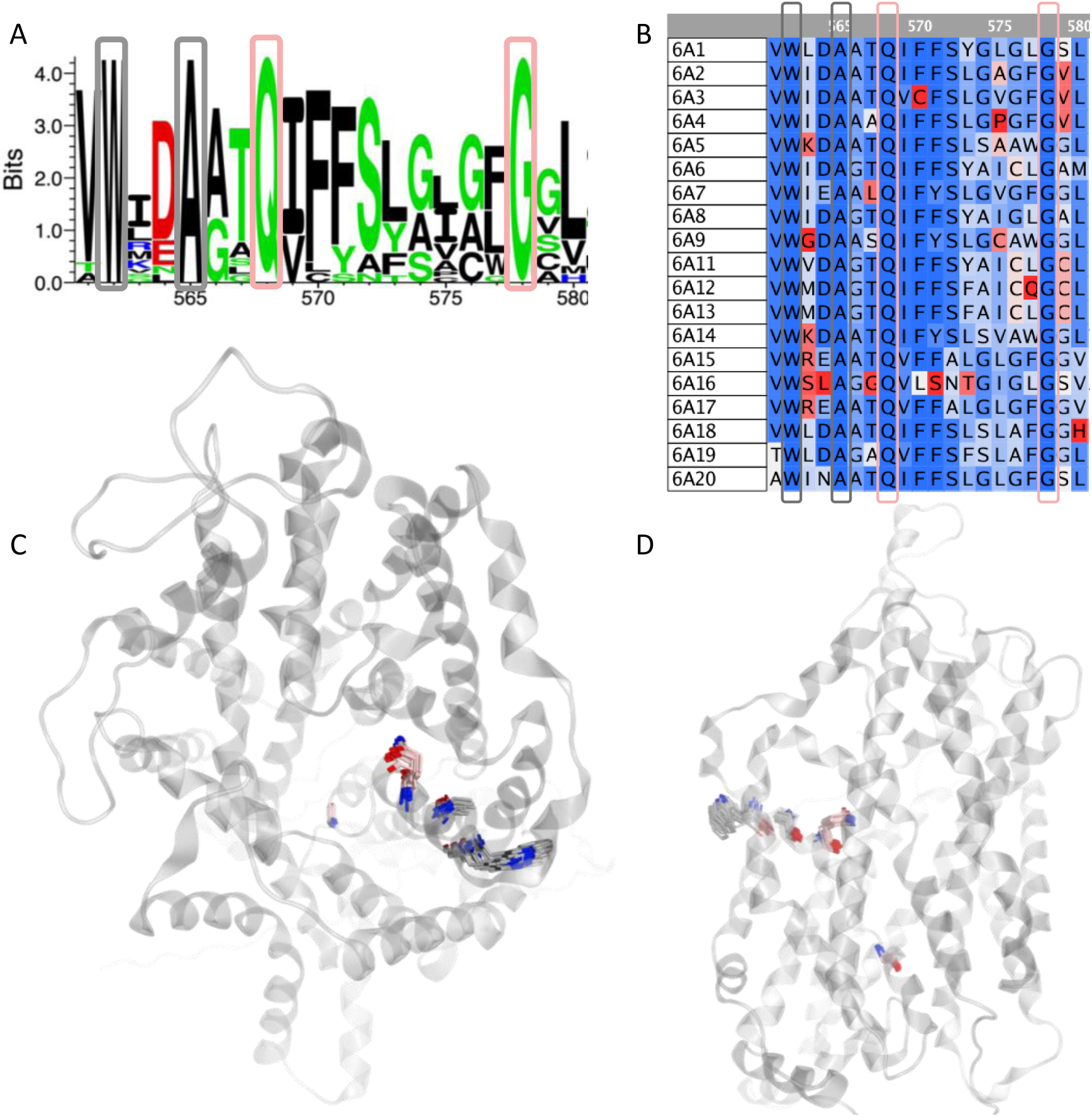
Sequence logo analysis of SLC6s canonical sequences. (A) TM6 fragment sequence logo plot, on which four residues were shown in full occupancy and conservation across the whole SLC6 family. Residues were colored with Tom Schneider’s color scheme. (B) TM6 multiple sequence alignment. Residues were colored based on their similarity. (C) The spatial location of the four residues from the top view. Gln and Gly (position on the alignment: 568 and 578) were presented in pink sticks, while the Trp and Ala (position on the alignment: 562 and 565) were in yellow. (D) The side view of the selected residue’s spatial location.

**Fig. 5.**
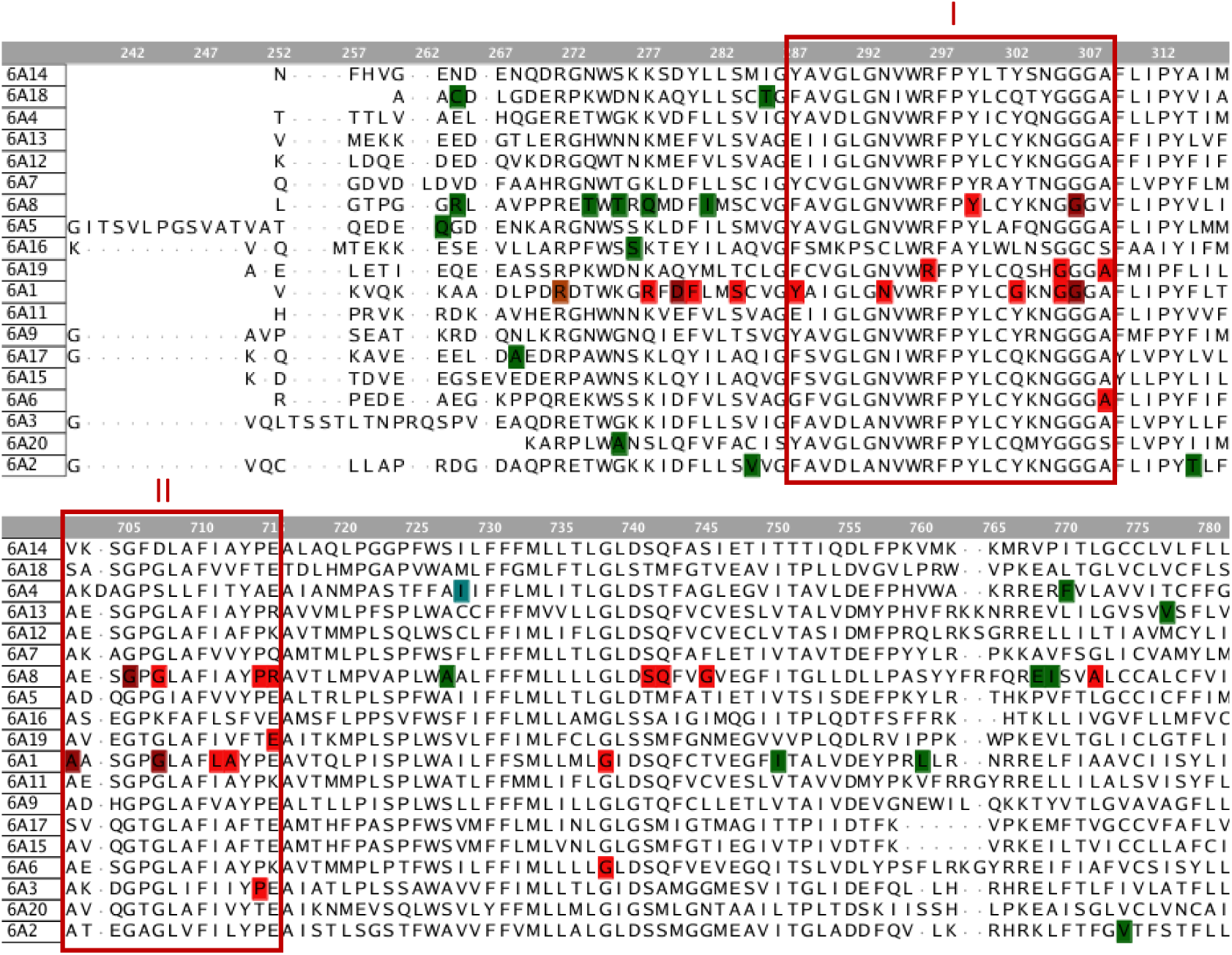
Distribution of mutation with clinical significance. Pathogenic mutations were colored in red and benign ones in green. Multiple mutation records on the exact residue location were indicated with deeper color shade. Two regions with explicit pathogenic mutation records were noticed and marked out with red boxes and annotated with I and II.

Compared to Gln and Gly on aligned positions 568 and 578, Trp and Ala on aligned positions 562 and 565 are less studied, despite all four residues being fully conserved across the SLC6s (Fig. 4B). When looking into the spatial location of these four residues (Fig. 4C), two observations can be made: One is that Trp and Ala sit less central but more peripheral than Gln and Gly. The other is that Trp and Ala adopted different orientations, with their sidechain pointing outside the transporter (Fig. 4D). These observations prompted us to consider the role of those residues in protein-protein interaction.

### Mutation spatial distribution

Upon conducting sequence alignment and structural superposition, signals of considerable conservation among the sequence, structure, and function emerged. To further explore this relationship in the SLC6 proteins, disease-related mutations offer a promising point of focus. From our previous work ^17^, a dataset of curated mutations with clinical significance labels was collected. These labeled mutations can serve as an entry point for analysis of their functional impact.

Two regions were identified with enrichment of pathogenic mutations. With the help of the mutation mapping tool (see methods), the spatial position of these two regions was located (Fig. 6). The region with aligned sequence numbers between 287 and 308 was found to be part of the transporting pathway. In such cases, mapping the region on the structure can already be informative for the pathogenicity of those mutations. However, in the case of the other region, the spatial location did not provide sufficient insight into the enrichment of pathogenic mutations in this region.

**Fig. 6.**
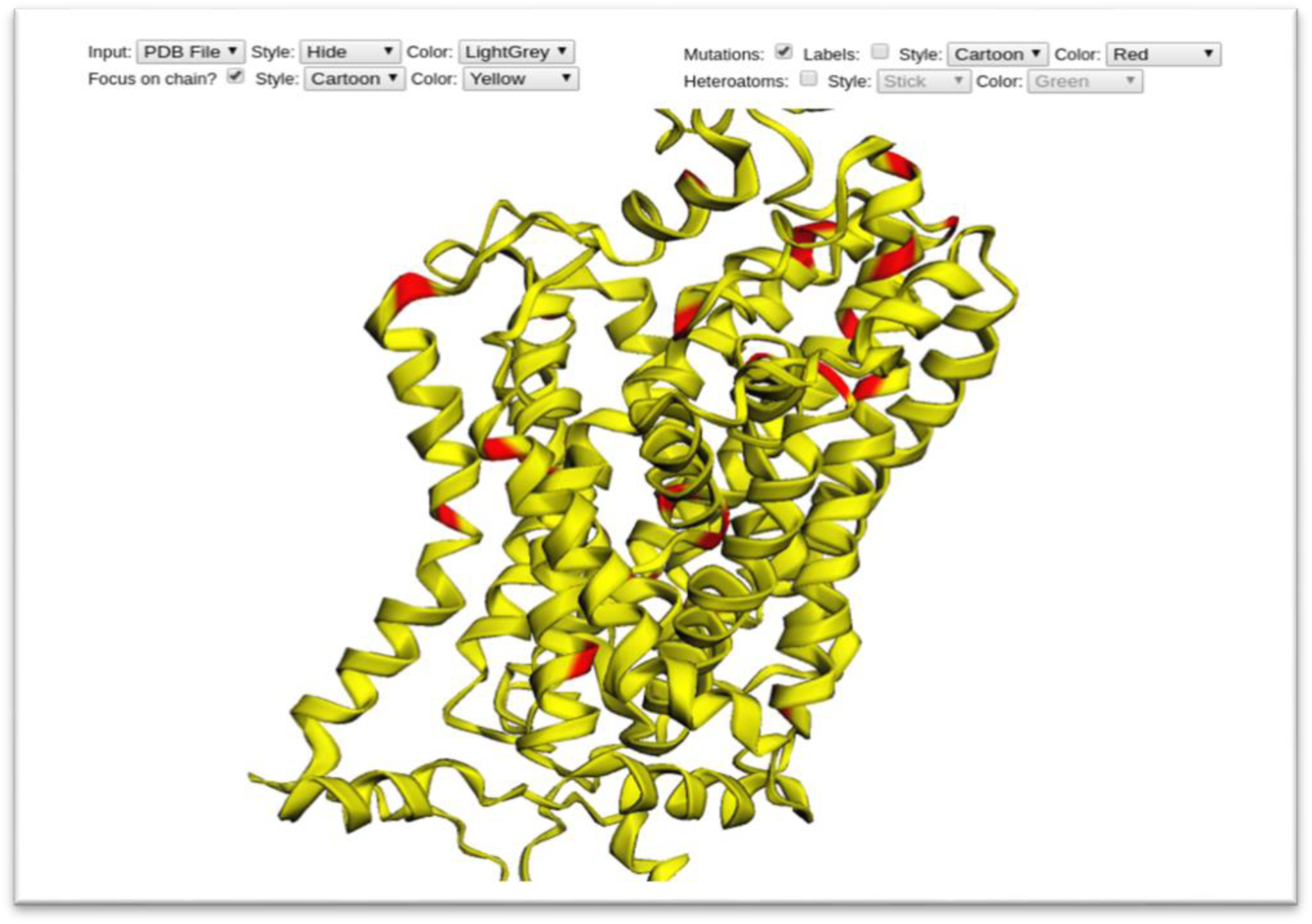
The mutation mapping tool display window. A set of SLC6A8 pathogenic mutations were displayed in this window. A region with enriched pathogenic mutations can be spotted on the extracellular surface located on the top right side of the window.

### SLC interactome extraction

The interaction proteomics dashboard can be found on the RESOLUTE website under the Resource section^23^. This release allows interactive network visualization for the defined individual SLC gene symbol and offers text data for downloading. Among the three available files on the website, the “SLC_interactome.txt” file was selected, which contains prefiltered interactions specifically curated to remove background and nonspecific protein-SLC interactions.

For further analysis, the interactome data was replotted with an automated Hotspot script (https://github.com/JiahuiH/HotSpots.git) integrating the following functions: (1) Filter the interaction data with a focus on the SLC6 family genes, (2) real-time adjust the stringency of the data shown by sliding the PPI probability threshold, (3) summarize the result for each SLC6 with the mean and sum metrics of filtered interactor PPI probability, and (4) generate an HTML link for the interactive network plot with the defined PPI probability.

While tuning the network plot, the clustering with shared interactors varied in size and the number of common interactors. In Fig. 7, an example is shown with a PPI probability threshold 0.9 for the whole SLC6 family. Seven out of 19 proteins were clustered in the biggest group. In this example, 16 common interactors were identified, with only one (SURF4) being a triple interactor. It needs to be mentioned that this visualization approach of selecting common interactors and interaction pairs is highly dependent on the defined threshold.

**Fig. 7.**
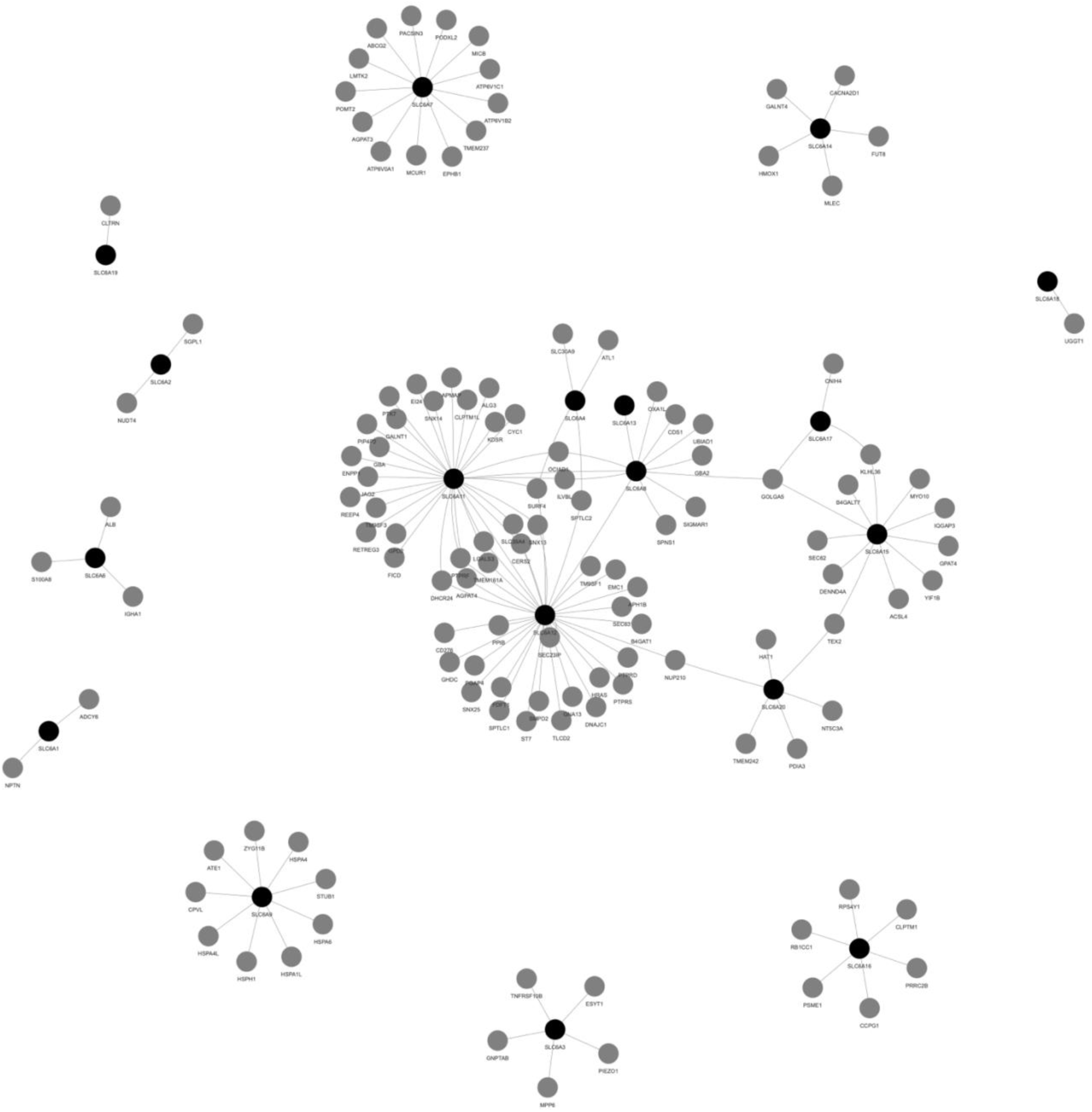
Network view of the SLC6s interactions. Black spheres represent the SLC6 proteins, and the interactive partners are shown in grey spheres with strings indicating the paired interactions.

To tackle this, a range of thresholds was applied, spanning from 0.6 to 0.9. This approach preserves the higher threshold to ensure precise, high-confidence interactions while also incorporating the lower threshold to capture a broader set of interactions. More precisely, the interactor was selected following these steps: Firstly, four lists of the interactors were collected with defined threshold values of 0.6, 0.7, 0.8, and 0.9. Secondly, each list was sorted according to the count of interactions. The minimum of count must not be less than two. Thirdly, the interactor present in the first order of all four lists was extracted. This ends up with one interactor, namely GOLGA5. Lastly, the consensus interactors within the second order of all four lists were included, which were SURF4 and OCIAD1. With this narrowed-down selection of the interactors (tab. 1), the pair of interactions for these three interactors was less strictly filtered in order to include as many SLC6 family members as possible. This was done by taking the interaction with a PPI probability higher than 0.6 with one of these three interactors.

**Tab. 1.** The final selection of interactors and their paired SLC6 proteins recorded in the SLCinteractome.

| Interactors | Paired SLC6s |
| --- | --- |
| GOLGA5 | SLC6A12, SLC6A15, SLC6A17, SLC6A7, SLC6A8 |
| SURF4 | SLC6A11, SLC6A12, SLC6A20, SLC6A4 |
| OCIAD1 | SLC6A11, SLC6A12, SLC6A4, SLC6A8 |

This selection covered 8 of 19 SLC6s, with at least one protein from one of the four subgroups of SLC6 (Nutrient amino acid transporters: SLC6A20, SLC6A15, SLC6A17; Neurotransmitter amino acid transporters: SLC6A7, Monoamine neurotransmitter transporters: SLC6A4; GABA, neurotransmitter, osmolyte, creatine transporters: SLC6A8, SLC6A11, SLC6A12). In total, 13 pairs of interactions were outlined for the subsequent procedures.

### Protein complex modeling and scoring

Thirteen paired interactions were extracted from the SLCinteractome data. These interactions were modeled with AF-M (V2.3) on the Life Science Compute Cluster. Each input file comprised two canonical sequences – the interactor and the paired SLC6 - with a total of about 1000 amino acids. Trough SLURM requests, eight CPUs, one GPU core, and 20G memory were assigned to each job. The time limit was set as 16 hours per job, which was sufficient to finish one job with the given computer power.

As output, 25 relaxed models, as well as the un-relaxed models, were provided by AF-M. Alongside a JSON format text file recorded the ranking of the relaxed models. This intrinsic model accuracy is determined with a weighted combination of pTM and ipTM scores described in the method section under “Model Evaluation Metrics”. In addition to the full model estimation indicated by the ranking score, pLDDT and predicted interface pLDDT scores were considered in this project to obtain a residue-wise evaluation of the predicted complex structure. Via scaled rank transformation, both scores were converted into the range of zero to one.

A family-wise enrichment of interaction positions was collected and plotted in two ways, as shown in Fig. 8. The collection was conducted by screening each modeled structure and extracting the residues in the complex interface. Predicted interface residues were captured by the modified pdb_paser script if any atom of the SLC6 protein residue was found within 3.5 Å of another atom from the paired protein. Those positive captured residues were mapped back to the aligned position with the scores the model possessed. When a relative position is well represented in the modeled complex structures, multiple sets of scores were processed as either mean or sum value, as shown in Fig. 8.

**Fig. 8.**
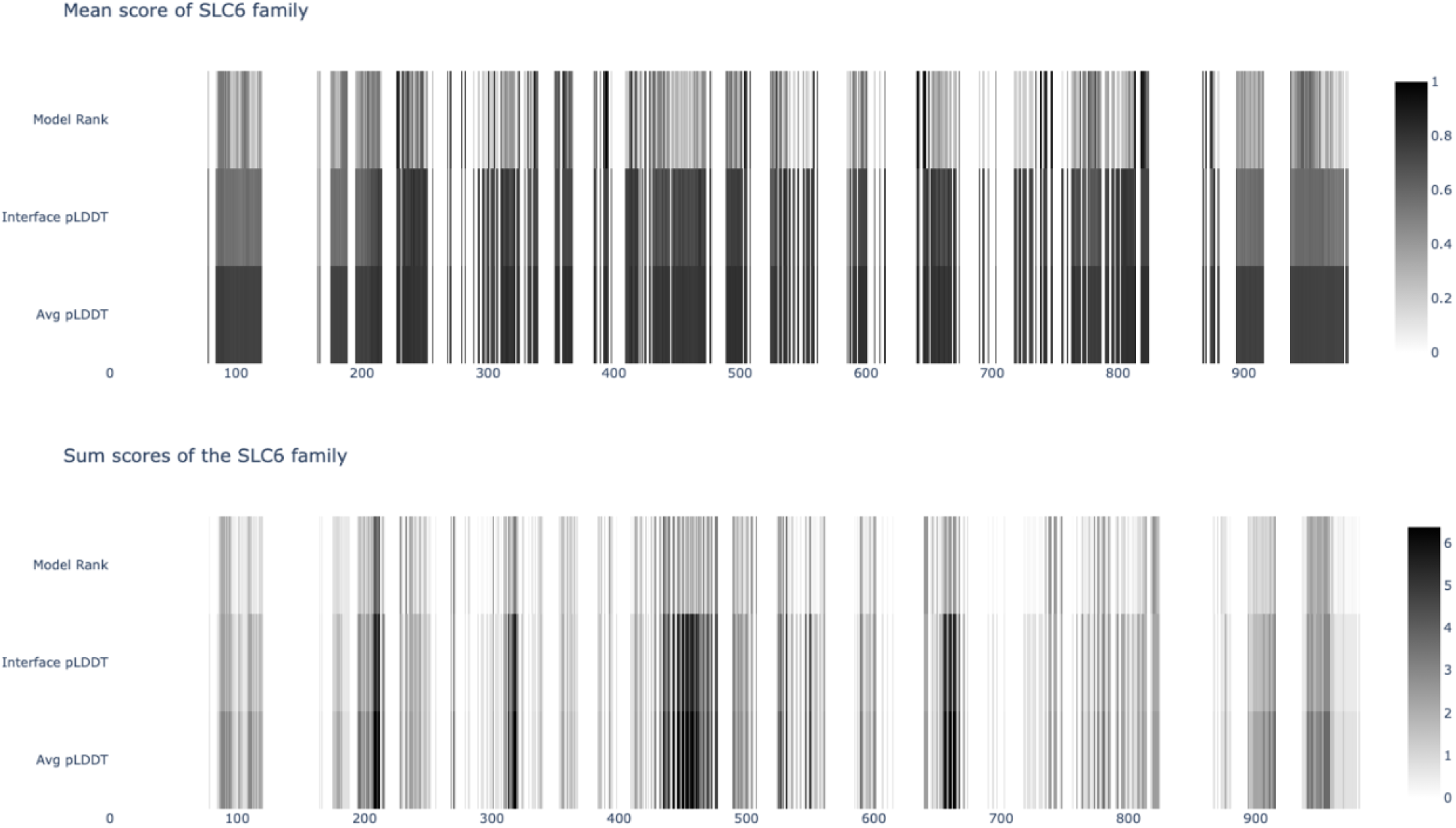
Family-wise enrichment of interaction positions. (A) The scores were averaged as mean values. (B) The scores were summed up on each aligned position. The scores plotted here were the model ranking, the interface pLDDT score, and the whole model averaged the pLDDT score.

Indicted by the protein interaction pattern generated with AF-M, these modelled complexes indicate an enrichment of key residues engaged in the PPIs. Compared to ligand-binding, PPIs are often characterized by a larger contact area, making it challenging for small molecules to fully occupy ^24^. Despite the relatively large surface of possible interactions, as suggested by the mean value of the scores, the sum values demonstrated more differentiable cavities of key interactive residues. Lu et al. ^25^ addressed this observation - termed as “hotspots” – explaining that those key residues are the core to stabilize the complexes. Mutation of even a single residue can disrupt complex formation, potentially leading to pathogenic consequences^26^.

To gain a first impression of the hotspot regions, the top-ranked models from each complex pair were extracted and mapped back to the paired SLC6 based on the aforementioned SLC6 superposition (Fig. 9). For clarity, undefined termini are excluded from the visualization. From this limited set of models, a preferred interaction pattern can be observed. For a more in-depth analysis, those regions were investigated in pairs of interactions.

**Fig. 9.**
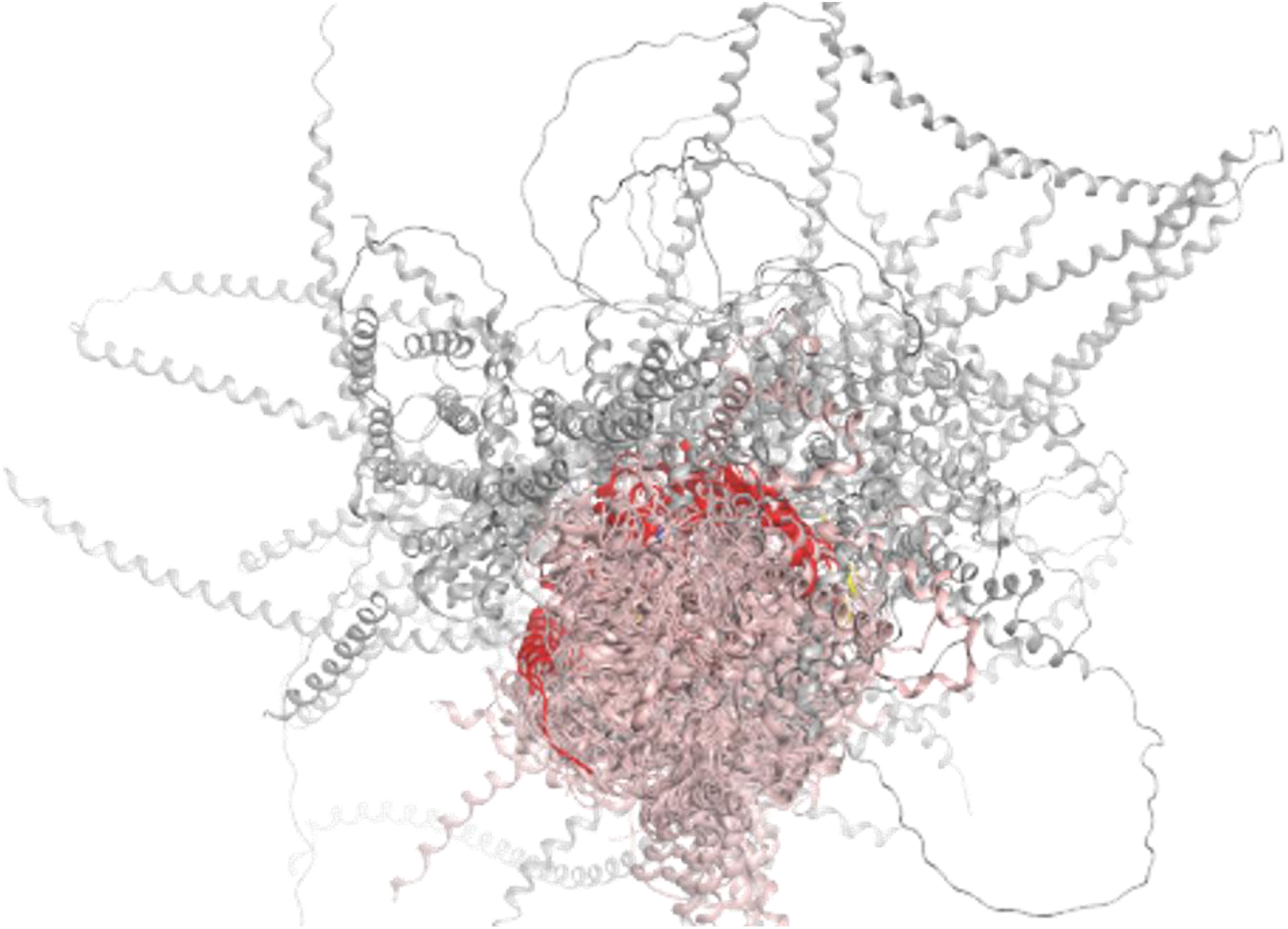
A top view of the top-ranked models from each modelled complex. All models were arranged according to the superposition of the SLC6 proteins in the complex. SLC6s are represented as pink ribbons, whereas the paired interactors are in grey. Two PPI hotspots are marked out in red.

### PPI with located mutation

The family-wise interaction hotspots highlighted regions with either mean score values indicating the characteristic protein-protein planar surficial interaction pattern or sum values pinpointing the key interactive residues. With this family-wise overview, an SLC6-shared PPI schema emerged. To further materialize the network, the hotspots captured in the family-wise analysis were expanded for more detailed investigations. Detailed interaction data were extracted from all modeled complexes. On top of that, the curated mutations with clinical significance were also incorporated into the analysis based on the hypothesis that mutations located in crucial PPI hotspots can often lead to pathogenic outcomes.

To gather insights from the disease-related mutations and the SLC6 PPI hotspots, both data sets were plotted together along the aligned full length of the SLC6 sequences (Fig. 10). For the modeled PPI, the interaction positions were first grouped based on the paired SLC6 and then mapped back to the aligned position. The score of the interaction depicted in this case for a more concentrated and rigorous study of the interface was the mean interface pLDDT score. More precisely, the elaborated interaction heatmap was displayed on the phylogenetic order of the SLC6 family, retaining the sub-family relationship of SLC6s. Moreover, the un-modeled SLC6 members were shaded to avoid confusion between non-binding residues and not-experimented sequences. The number of pathogenic mutations from the previous work ^17^ was counted at each aligned residue position and plotted on top of the interaction heatmap.

**Fig. 10.**
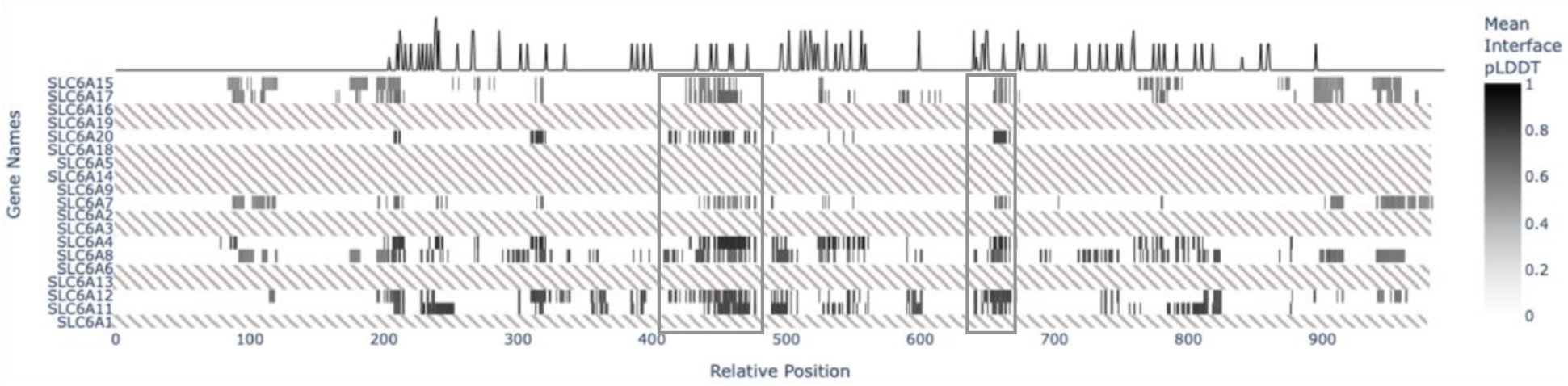
Heatmap of detailed interaction data from all modeled complexes together with the line plot of the count of curated pathogenic mutation displayed on the aligned full length of SLC6 sequences. The two grey boxes mark out two regions where PPIs were captured for all modeled complexes, and mutation records were also clustered.

One of the regions depicted well from all modeled complexes was mapped back to a fragment spanning from helix 5 to helix 6 (also known as ECL3, Fig. 12A). Signals of pathogenic mutations were also present in this region, although not recorded on the SLC6 members that were selected for modeling. As compensation for the limited collection of disease-related mutations, we streamlined the AlphaMissense database and observed a consented pattern of the pathogenicity probability in this region. From the predicted pathogenicity probability plot, an intriguing highly pathogenic residue (marked out with yellow box in Fig. 11) was detected next to this region. It was recognized as a fully conserved proline adjacent to the ECL3 to allow the flexibility of the extracellular loop before the start of helix 6. A paired interaction between SURF4 and SLC6A4 is illustrated in Fig. 12B. On the SLC side, two somatic mutations were detected in a large-scale cancer study ^27^, namely Leu310Ppro (Genomic Mutation Identifier: COSV55568226) and Leu313Ser (Genomic Mutation Identifier: COSV55570177). On the other side of this hotspot, multiple non-somatic mutations on SURF4 residue Trp45 and Phe71 can be found on Ensembl ^28^ and are annotated as deleterious. For instance, Trp45Cys (Ensembl: ENST00000371989), Phe71Leu (Ensembl: ENST00000371989), and Phe71Ser (Ensembl: ENST00000629578).

**Fig. 11.**
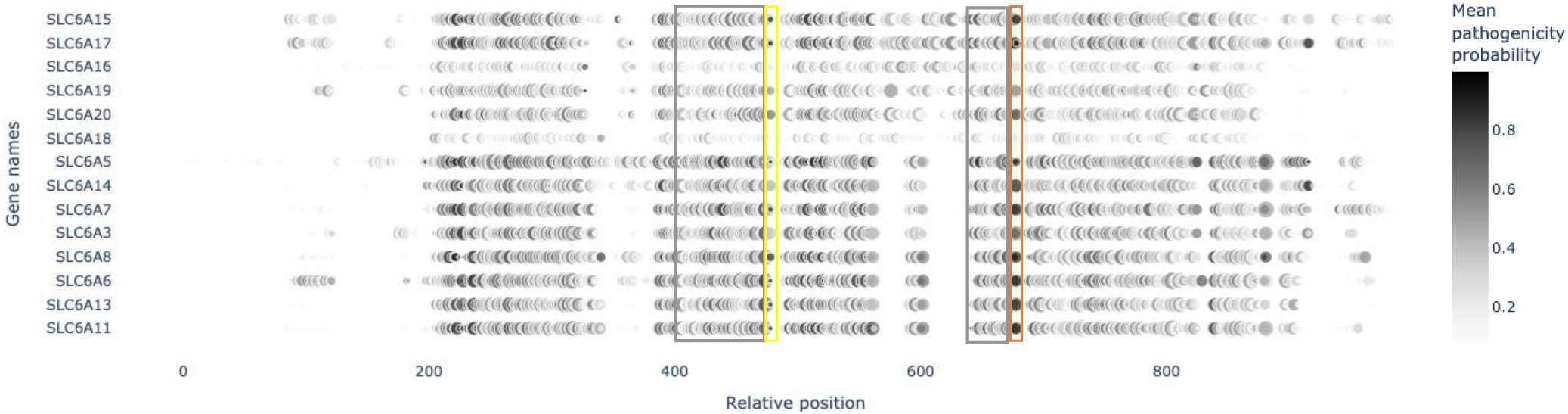
AlphaMissense prediction of SLC6s mutation pathogenicity probability. The value on each aligned position was calculated as the mean of all possible variants’ pathogenicity probability, with the circle radius indicating the standard deviation of the values. Grey boxes correspond to the two PPI hotspots outlined in Fig. 10. The Yellow box highlights a highly pathogenic residue next to one of the hotspots. The orange box marks out the fist residue of helix 8 adjacent to the other hotspot.

**Fig. 12.**
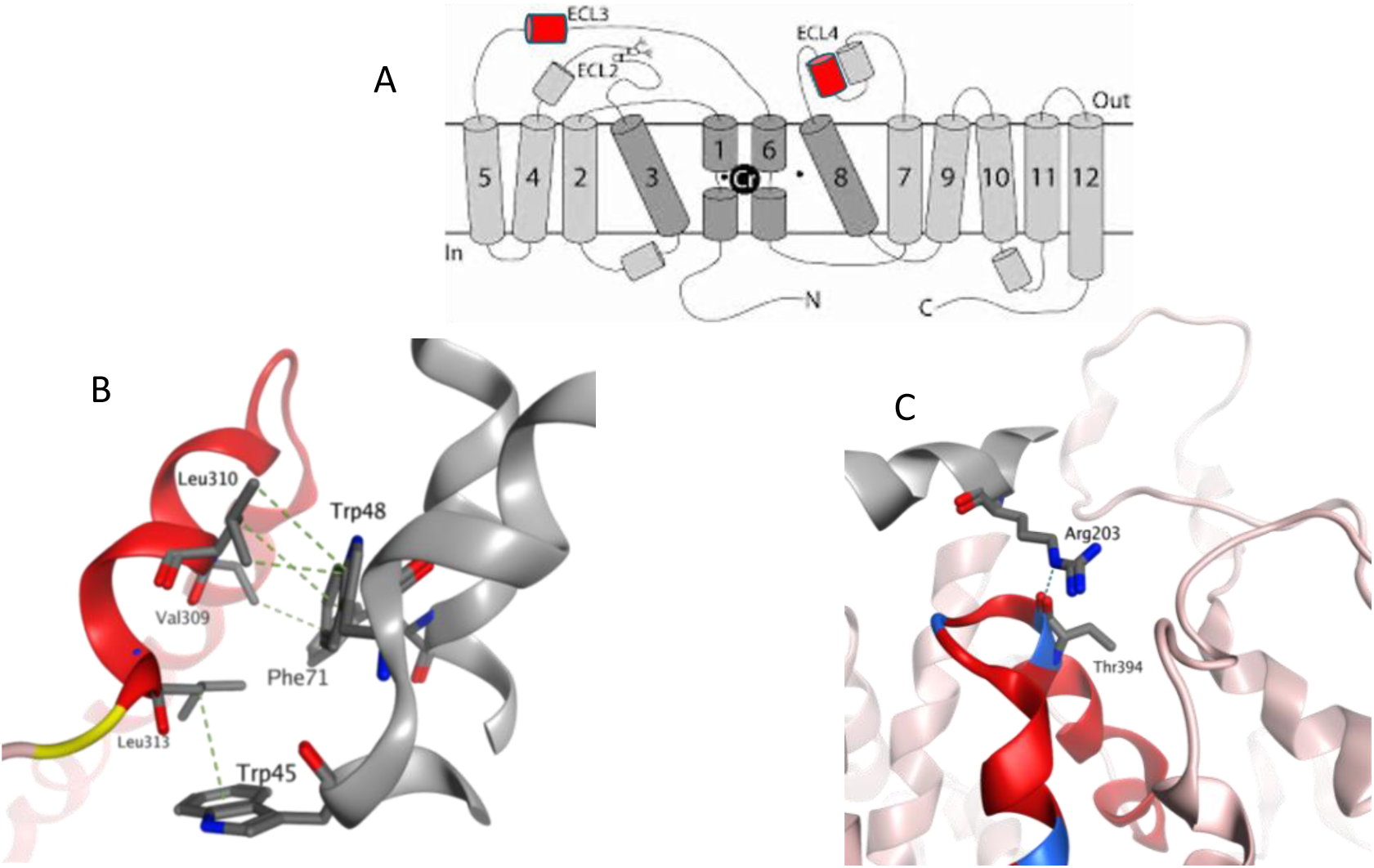
Two PPI hotspots and examples of modeled interaction in these regions. (A) Schematic view of the SLC6 topology with cylinder indicating the defined secondary structure. Two Hotspots are marked out in red. Figure adapted from Salomons et al. ^35^ (B) One hotspot corresponding to ECL3 demonstrated a tight interaction in the modeled SURF4-SLC6A4 complex. (B) Another hotspot corresponding to ECL4 exhibited an interaction between the modeled OCIAD1-SLC6A8 complex. H-Pi interaction is shown in green dashes and H-bond in blue.

The other region marked out with a grey box in Fig. 11 was tracked on the extracellular surface, which was previously noticed as an enriched region of pathogenic mutations with the mutation mapping tool. This region mainly comprised a fragment of ECL4 and the loop connecting ECL4 to helix 8 (Fig. 12A). The biological role of helix 8 in SC6s is well-recognized as part of the substrate-transporting pathway. Correspondently, next to this hotspot, an unneglectable high signal of mutation pathogenic probability can be observed from the AlphaMissense prediction as annotated with the orange box in Fig. 11. Located adjacent to helix 8, this hotspot can be recognized by a series of crucial structural motifs. ECL4 was reported in various publications ^29–31^ functioning as part of the lid shielding the pathway from the extracellular solvent. To fulfill this function, both defined helical packing and the flexible loop are required so that the lid can fold against other helices in the bundle and form stable interactions during conformational change ^32^. Although the current knowledge regarding the structural composition of this hotspot primarily emphasizes its role in conformational changes, there is also evidence of mutations in this region causing significant enrichment of ER protein partners (Thr394Lys, Pro397Leu, and Ala404Pro in SLC6A8 ^33^). One of our modeled complexes provided an illustration of possible interactions between OCIAD1 and SLC6A8 (Fig. 12C). OCIAD1 is an endosomal protein crucial for cell potency ^34^. The interactive residue Arg203 on OCIAD1 was annotated on Ensembl with two deleterious missense mutations (Arg203Lys and Arg203Met).

## Discussion and Conclusion

This work provides a perspective to study the impact of mutations within specific regions of the SLC6 family on the protein-protein interaction. Through multiple sequence alignment (MSA) and fine-tuned structural superposition, we quantify the degree of conservation in SLC6 proteins^36^ with specific pair-wise similarity values, highlighting not only their sequence-level conservation but also a shared structural pattern. Further investigations were conducted on both levels to exemplify the relationship between transporter function and this conserved pattern. On the sequence level, the sequence logo analysis revealed more details about the conservation of key residues. For example, the highly conserved Gln and Gly in TM6 play crucial roles in the function of several SLC6 members, such as their involvement in chloride ion binding and protein dimerization. Interestingly, some conserved residues located peripherally, such as Trp and Ala, were comparably less studied. On the structure level, through mapping the mutations with clinical significance on the structure, regions with enriched pathogenic mutations emerged. Two regions were highlighted at this stage: one within the TM helices and the other in the extracellular loop (ECL) regions.

Further exploration was performed from the perspective of protein-protein interactions regarding the highly conserved residues located peripherally observed from the sequence level and the enriched pathogenic mutations on the extracellular surface spotted from the structure level. The SLC interactome data was utilized for the extraction of SLC6 interactors, which were selected as GOLGA5, SURF4, and OCIAD1. In total, 13 paired complexes were modeled using AlphaFold-Multimer, highlighting two key hotspots on the SLC6 proteins. One referred to ECL3, and the other to ECL4. These regions showed enriched interactions with interactors and contained several pathogenic mutations. As compensation for the limited collection of mutation data with clinical significance labels, AlphaMissense prediction was integrated. It offered further support to the significance of these hotspots. The modeled OCIAD1-SLC6A complex exhibited the PPI on the ECL4 hotspot, offering a hypothetical illustration for the deleterious effects of the mutations regarding their subcellular localization findings.

In conclusion, this study demonstrated the importance of conserved structural features and interaction hotspots in the SLC6 family. The identification of PPI hotspots, along with the pathogenic mutations located within them, suggested the important role of these regions in both the structural integrity of the SLC6 transporters and their interactions with other proteins. This study also underscores the importance of large-scale investigations led by the RESOLUTE and REsolution consortia^23, 37–39^, which provided a crucial foundation for the computational modeling and predictions presented in this work. Nevertheless, this study also encounters certain limitations. The collection of clinically significant mutation data is limited, restricting our ability to comprehensively map out the region with clinical significance. Likewise, predictive models for mutation pathogenicity were opted in this case as compensation. Furthermore, not all SLC6 family members were modeled with the selected interactors, which could result in an incomplete understanding of the full range of potential interactions and structural adaptations across the family. Addressing these limitations in future research could provide more robust, experimentally validated insights into the impact of mutations on the protein-protein interaction of the SLC6 family.

Overall, the integration of sequence, structure, protein-protein interaction, and mutation data provided a comprehensive view of how conserved regions and disease-related mutations contribute to the understanding of both normal function and pathogenic outcomes of these proteins.

## Methods

### Multiple Sequence Alignment

The Clustal Omega ^40^ algorithm was chosen to conduct the multiple sequence alignment (MSA) for its efficiency in computing large alignment for protein family with a high accuracy compared to alternative packages ^41^. An alignment file was exported from the EMBL-EBI tool service ^42^ for further operations. Colors cannot be saved as part of the alignment. It was then visualized and colored on molecular operation environment (MOE) with Clustal-X color scheme ^43^. A pairwise similarity matrix was applied for the interpretation of the alignment file. This matrix calculates the positive matches between sequence i and j divided by the sequence length of j. A positive match is taken when the paired residues have BLOSUM62 greater than zero.

The purpose of creating MSA is to gain a first impression of the mutation distribution from the sequence perspective. This step is not required for the structural modeling because the MSA from AF-M is pairwise to the interaction partner instead of across the SLC6 family.

**Table:** Colour scheme of mutation counts on the MSA.

|  | one record | two records | three and more records | one pathogenic and one benign |
| --- | --- | --- | --- | --- |
| Pathogenic | red | dark red | brown | magenta |
| Benign | green | cyan | blue | magenta |

### Sequence logo

To better understand the SLC6 MSA, a sequence logo ^44^ was generated with the alignment CLUSTAL file. It can provide a graphic view of the sequence conservation, depicting the consensus residue while preserving the diversity of the sequences. Not all tools available ^45^ for computing sequence logos are suitable for plotting the protein sequence. After checking availability, the tool Seq2Log ^46^ was chosen in consideration of accessibility and reasonable responding time.

The set-up on the Seq2Log web server was defined as the following: (1) Select logo type as Shannon entropy ^47^. (2) Weight on prior as 0. (3) Select information content units as bits. (4) Amino acids color scheme as Tom Schneider’s ^44^. Other than these settings, all the rest are kept as default.

The logo type is specified as Shannon entropy instead of the other relative entropy options (Kullback-Leibler, Weighted Kullback-Leibler, Probability Weighted Kullback-Leibler) because it measures the variability of a distribution rather than comparing two distributions. For a multiple protein sequence alignment, the Shannon entropy (H) for each residue position is calculated as follows:

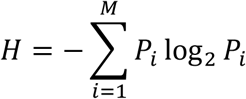

Where P_i_ represents the proportion of amino acid type i, and M refers to the total number of amino acid types (20 in total). The entropy value H can range from 0 (indicating only one amino acid is present at that position) to 4.322 (indicating all 20 amino acids are equally represented at that position). Generally, positions with H>2.0 are considered variable, while those with H<2.0 are deemed conserved. Positions with H<1.0 are considered highly conserved ^48^.

Weight on prior was set as 0 in order to decline the pseudo count correction for low counts that Seq2Logo includes as default. With the default feature, the amino acid frequencies displayed in the sequence logos are corrected for low number of observations using a BLOSUM (BLOcks SUbstitution Matrix) amino acid similarity matrix ^49^.

Lastly, the information content unit (y-axis) is not displayed as the entropy but as the value of the bits. It measures the reduction in uncertainty of a given sequence alignment compared to a random one. This is calculated by subtracting the entropy of specific amino acids at a position, H, from the entropy of the position if the alignment were random.

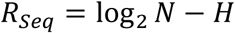

In protein sequence alignments, log_2_ *N* is always log_2_(20) = 4.32. Therefore, if only one residue is present at a given position, the R_seq_ will be 4.32, as H equals 0, indicating very low variability at that position.

### Mutation mapping tool

To get a fast and automatic mapping of a list of mutations onto the 3D structure of the respective transporter, a mutation mapping tool was compiled on the open-source data analytics platform KNIME ^50^. This tool was built by integrating the publicly available JavaScript library 3Dmol.js ^51^. It utilized the Web Graphics Library (WebGL) to accelerate the rendering of interactive 2D and 3D graphical operations on web browsers without any additional plugins ^52^.

As input, it takes tabular formats of listed mutations. Preferably, defining a column containing annotations to differentiate the mutations can be beneficial for further analysis. As for the protein structure, both PDB and the AlphaFold_DB are integrated into the options with a given PDB or UniProt Identifier. The last step before the mapping is a position checking to avoid mismatching of isoform sequence mutations being mapped on the structure models based on the canonical sequences.

### SLCinteractome

We retrieved the protein-protein interaction information from the SLC interaction proteomics database released by RESOLUTE ^23^. The dataset contains 1747 mass spectrometry injections and a total of 3.2 TB of raw data. It surveyed 396 SLCs by means of affinity purification coupled with quantitative MS(AP-MS). This work was conducted under the framework of the RESOLUTE and REsolution consortia, applying a systematic omics approach to enrich the protein interaction data on SLCs.

The current interactome data are limited in their information on SLCs, as they do not employ tailored protocols for transmembrane-spanning proteins. When compared to isolated interactome analysis of individual membrane proteins, this large-scale study eliminated the common cellular interaction modalities using AP-MS. Over 400 SLCs were purified from human embryonic kidney cells. After cross-checking with available individual SLC-interactiomes and screening with a machine learning method, a total of 18,991 interactions for 396 SLC proteins across 68 different SLC families were retrieved.

Their keynote to separate significant interactions from the background was a combination of two strategies: One was to set a combined scoring feature with ComPASS ^53^ and SAINTexpress ^54^. The other was to derive quantitative features such as summed mean spectral counts. An ensembled Radial Basis Function (RBF) based classifier was fitted to the aforementioned features training on a data set with 693 true and 970 false labels.

### AlphaFold Protein Structure Database

Initially released in 2021, the AlphaFold Protein Structure Database (AF-DB) expanded from 365 thousand to 214 million predictions with its latest release ^19^.The whole SLC6 family structural models were extracted from the latest release for the benefit of full-length predictions of the canonical sequences. The extraction was performed with the UniProt identifier of the canonical SLC6 sequences.

### AlphaMissense

DeepMind released in 2023 their new AI model AlphaMissense ^55^ in the application of the missense variant classification, together with a database covering 89% of all 71 million possible missense variants. The main architecture of the model is based on the AlphaFold2 framework with fine-tuning on the labeled human or close-homologue variants data. As output, the missense mutations will be categorized as pathogenic, ambiguous, or benign, along with a pathogenicity probability. Predictions for all human missense variants with the file name AlphaMissense_hg38.tsv.gz was downloaded from the correspondent Zendo depository ^56^.

The data was processed for the purpose of this study as follows: The loaded data was converted from a TSV file into a pandas Data Frame in chunks, extracting crucial information such as key transcript ID, protein variant annotation, pathogenicity class, and pathogenicity probability score. The extracted data was aggregated by grouping the transcript ID and variant information and later saved as an aggregated CSV file for future use. Subsequently, the aggregated data was filtered to match relevant transcript IDs from the sequence alignment. Based on the alignment, the mutation position annotations were converted to relative positions. To condense the information for each sequence and enable the visualization of the pathogenicity probability prediction across the whole family, the mean and standard deviation values were calculated.

### AlphaFold-Multimer

The AlphaFold-Multimer is built on top of the AlphaFold2 architecture. It extends from the application domain of AlphaFold2 to predict multi-chain proteins. This functionality was applied to the protein complex structure modeling ^57^. Compared to AlphaFold2, AF-M introduces a novel way of selecting subsets of residues for training, such that the model learns more about intra-chain interactions. It can be applied by defining the model_preset flag of the command as multimer instead of monomer. The max_template_date flag should be carefully set when there are experimentally solved structures that should be excluded from the modeling template set consideration (In this work: --max_template_date=2020-05-14). To maintain the accuracy reported in the AF-M paper, the seed number was kept at 5. In total, 25 predictions were generated per input.

Another requisite is an input fasta with multiple sequences. In each fasta file, there should be two sequences. One is the prey or proposed interacting partner sequence, and the other is the bait or target protein sequence. Each sequence should contain no line breakers and start with >UniprotID. In this way, the output file will be generated and named in a human-interpretable way.

The modeling process demands significant computational resources due to its complex architecture and the need to access the full AlphaFold genetic databases, which in total reach 3 TB. Therefore, we pre-processed the FASTA file on a local machine and later submitted the job to the Life Science Compute Cluster (LiSC), run jointly by the University of Vienna and the Medical University of Vienna.

### Model Evaluation Metrics

The resulting relaxed models of the AlphaFold-Multimer are named by default with the model confidence, determined by the following formula:

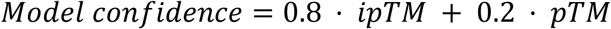

Predicted TM score (pTM) is a metric inherited from the AF. Nonetheless, as a protein complex modeling approach, AF-M signed higher weight to the interface predicted TM score (ipTM). The pTM score provides one score per structure, estimating the structural similarity between the predicted model and the true structure. Another structure quality metric also originated from AF — the predicted local distance difference test (pLDDT) score — which offers a more detailed structure evaluation by measuring the confidence for each individual residue.

The predicted interface pLDDT score was calculated with a script adapted from the “pdb_parser” published by Andrea Rubbi on Github ^58^. This script detects the atoms from two chains of the complex that are within a certain threshold. Besides heavy atoms, also hydrogen is included in the atom consideration. The threshold was set as 3.5 Å. The code was modified in the following perspectives for a better fitting to the task and higher efficiency: (1) the output file was renamed according to the input to prevent output overwriting, (2) the heme detection part of the original code is removed as it is unrelated to our task and hinders the processing speed, (3) the printing graph was simplified to show only when each output was generated, and (4) the inquiry of the chain was moved out from the input function into command line arguments so that it could be defined in another script. On top of this, another script, loop_parser.py, was created to loop over all run results and conduct the modified pdb_parser for each PDB file. This loop was parallelized to fully utilize all the available threads. Because of the looping, every PDB file will have a CSV file generated by the pdb_parser. The CSV file will contain the following information: The potential iterating partner on chain A, the residue number, and the atom type. The same information will be recorded for the chain B side, as well as the distance between these two atoms from two chain.

Noting that the pLDDT score ranges from 0 to 1 and the model ranking score ranges from 0 to 24, these scores are not directly comparable due to their differing scales. Additionally, since the 25 models were generated using random seeds, there is no inherent assumption of linear improvement across the models. To address this, a scaled rank transformation was applied. This transformation ensures that both sets of scores are comparable while preserving the original distributional characteristics of each set. For a collection of scores the X = {x_1_, x_2_,…, x_i_}, the scale rank of each value x_i_ is calculated as:

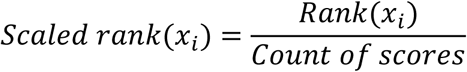

Where the count of scores for both collections of scores equals 25, as the AF-M generates 25 for each job.

## Acknowledgements

This work was performed within the REsolution project. RE*solution* has received funding from the Innovative Medicines Initiative 2 Joint Undertaking (https://ihi.europa.eu) under grant agreement no. 101034439. This Joint Undertaking receives support from the European Union’s Horizon 2020 Research and Innovation Programme and EFPIA. This article reflects only the authors’ views and neither IMI nor the European Union and EFPIA are responsible for any use that may be made of the information contained therein. This work was also supported by the Austrian Science Fund/FWF, grant W1232 (MolTag). This work utilized the computational resources of the Life Science Compute Cluster (LiSC). We also thank Eva Hellsberg for her feedback on the manuscript.

